# Dissociation in basolateral and central amygdala effective connectivity predicts the stability of emotion-related impulsivity in adolescents with borderline personality symptoms: a resting-state fMRI study

**DOI:** 10.1101/2021.05.17.444525

**Authors:** Nathan T. Hall, Michael N. Hallquist

**Affiliations:** Department of Psychology and Neuroscience, University of North Carolina at Chapel Hill

**Keywords:** borderline personality, adolescence, effective connectivity, resting-state, amygdala, urgency, emotion-related impulsivity

## Abstract

**Background:** Borderline personality disorder (BPD) is associated with altered activity in the prefrontal cortex (PFC) and amygdala, yet no studies have examined fronto-limbic circuitry in borderline adolescents. Here, we examined the contribution of fronto-limbic connectivity to the longitudinal stability of emotion-related impulsivity (ERI), a key feature of BPD, in symptomatic adolescents and young adults.

**Methods:** We compared resting-state effective connectivity (EC) in 82 adolescents and emerging adults with and without clinically significant borderline symptoms (n BPD = 40, ages 13-30). Group-specific directed networks were estimated amongst fronto-limbic nodes including PFC, ventral striatum (VS), central amygdala (CeN), and basolateral amygdala (BLA). We calculated directed centrality metrics and examined if these values were associated with initial levels and rates of change in ERI symptoms over a one-year follow-up using latent growth curve models (LGCMs).

**Results:** In the healthy group, ventromedial prefrontal cortex (vmPFC) and dorsal ACC had a directed influence on CeN and VS respectively. In the borderline group bilateral BLA had a directed influence on CeN, whereas in the healthy group CeN influenced BLA. LGCMs revealed that in borderline adolescents, ERI remained stable across follow-ups. Further, higher output of R CeN in controls was associated with stronger within-person decreases in ERI.

**Conclusions:** Functional inputs from BLA and vmPFC appear to play competing roles in influencing CeN activity. In borderline adolescents BLA may predominate over CeN activity, while in controls the ability of CeN to conversely influence BLA activity is associated with more rapid reductions in ERI.

---

Borderline Personality Disorder (BPD) is characterized by affective instability, interpersonal dysfunction, suicidality, and self-harming behaviors (1). Although BPD is most often diagnosed in emerging adults, there is mounting evidence that symptoms often begin in adolescence and show continuity into adulthood (2–5). This developmental course aligns with broader research on impulsivity and emotion regulation in adolescence (6). In BPD, impulsive and self-destructive behaviors tend to occur in response to momentary emotional arousal and constitute a core symptom of the disorder (4,7,8).

Difficulty controlling impulses in the face of negative emotions (e.g. self-injury after a break-up with a romantic partner) is referred to as Negative Urgency (NU) (9,10). Positive Urgency (PU) is a related feature (11,12) that describes a tendency to act impulsively in order to enhance positive mood (e.g. engaging in risky sexual behavior with a stranger while intoxicated). Collectively, NU and PU are referred to as emotion-related impulsivity (ERI) and reflect a crucial intersection of emotion regulation capacities and impulsivity that is robustly associated with psychopathology (13). At the level of neural circuits, ERI may reflect an imbalance between the inhibitory control functions of ventromedial prefrontal and orbitofrontal cortex (vmPFC, OFC) and emotion-congruent response tendencies in the amygdala and ventral striatum (VS) (13,14). For example, even mild uncontrollable stressors can profoundly disrupt PFC functioning (15), leading to a reliance on short-sighted, emotion-congruent behaviors that are immediately reinforcing, to the detriment of long-term wellbeing and safety.

Neuroimaging studies of emotion in adults with BPD consistently identify abnormalities in similar fronto-limbic circuits, including the amygdala, VS, medial PFC, and dorsal anterior cingulate cortex (dACC) (16–21). More specifically, several studies have noted a fronto-limbic imbalance in BPD: in response to a range of emotional stimuli, limbic regions are often more active whereas activity in prefrontal regions involved in emotion regulation (esp. mPFC and ACC) is blunted (17,18,22–24). Fronto-limbic accounts of emotion dysregulation center on the amygdala, which is involved in detecting threat, representing the emotional significance of stimuli (25), and encoding and retrieving fear memories. Furthermore, emotional experience depends on interactions between the amygdala and mPFC and dACC, which are involved in emotional appraisal and regulation (26).

Preclinical research casts the amygdala within a larger circuit that includes the cortex, striatum, and midbrain. Together, this circuit is fundamental to the experience and expression of emotional behaviors (27). Further, nonhuman animal studies demonstrate that the central nucleus of the amygdala (CeN) and the basolateral amygdala (BLA) play dissociable roles in the generation of Pavlovian associations and retrieval of relevant unconditioned responses during instrumental learning (27,28). Broadly speaking, the phylogenetically old CeN is a major controller of the autonomic nervous system, having strong projections to hypothalamus, periaqueductal gray (PAG), basal forebrain, and brainstem — brain regions contributing to autonomic arousal (29). In contrast, the phylogenetically newer BLA receives direct inputs from sensory cortex and uses this information to construct an emotional representation of specific conditioned stimuli and uses this information to influence the output of CeN (27,30). This body of research suggests that the expression of emotion-congruent behaviors depends on BLA’s capacity to convey the emotional significance of specific conditioned stimuli to CeN, which projects to regions that affect arousal and initiate approach and avoidance behaviors (27).

Further, amygdala and PFC functionally couple with the ventral striatum (VS), forming fronto-striatal-limbic loops that support reward learning and motivation (31). Studies of reward learning consistently find activity in VS in anticipation of reward (32,33) and after the receipt of reward (31,34). In studies of impulsivity, impaired dopaminergic functioning in VS discounts the value of future rewards, tilting choices toward more immediate rewards (35–37). Extant research suggests that VS function is impaired in individuals with BPD (19,38,39), though a fine-grained analysis of VS contributions to fronto-limbic abnormalities has not been a focus of the BPD neuroimaging literature.

Connectivity of both amygdala and VS show pronounced developmental changes in adolescence that are susceptible to stress (40–42), yet these connections have not been investigated in adolescents with borderline symptoms. This is surprising given that the emergence of BPD symptoms is associated with trauma, interpersonal discord, and chronic stress (43,44). Importantly, during the transition from mid-adolescence to early adulthood, selfreported impulsivity shows marked mean-level decreases in the general population^1^ (45,46). However, within-person changes in impulsive symptoms are heterogenous. Some highly impulsive adolescents show relative stability in impulsivity or decrease only slightly during this period, potentially leaving these individuals vulnerable to persistent negative outcomes into their 20’s (46,47).

Altogether, while the fronto-limbic account has received attention in adults with BPD, little is known about fronto-limbic disturbances in adolescence, when the development of ERI may crucially impact the development and maintenance of BPD symptoms. In this resting-state fMRI study of adolescents and emerging adults with BPD symptoms, we examined fronto-limbic circuitry using effective connectivity (EC) analyses within a network neuroscience framework (48). Furthermore, we tested how fronto-limbic connectivity related to within-person stability and change in ERI over six- and twelve-month follow-up assessments. Motivated by evidence that effective connectivity of R CeN differed substantially between groups, we tested the hypothesis that R CeN connectivity underlies the relationship between clinical group membership (healthy control versus BPD) and within-person change in ERI symptoms. The results of the current study point to a dissociation in EC between R CeN and BLA that accounts for the stability of ERI in symptomatic individuals over the course of a year.

## Methods and Materials

### Participants

Participants were 46 adolescents and emerging adults with BPD symptoms recruited from community and outpatient settings, as well as 44 sex- and age-matched healthy controls. All participants were screened using the Personality Assessment Inventory-Borderline scale (49) with BPD participants screening ≥ 30 and controls screening < 17. The average age was 20.53 years (range 13-30 years); 59 participants were female and 31 were male. Eight participants (*n* BPD = 6) were excluded from our analyses for having poor fMRI data quality (Supplemental Methods). See Table 1 for a complete demographic characterization of the final sample.

**Table 1.** Sample Characteristics

| Characteristic | BPD (*n* = 40) | HC (*n* = 42) |
| --- | --- | --- |
| Age (SD) | 20.84 (4.42) | 20.61 (4.16) |
| PAI-BOR (SD) | 47.3 (9.60) | 10.3 (3.99) |
| Ethnicity |  |  |
| Hispanic or Latino | 3 | 2 |
| Not Hispanic or Latino | 36 | 40 |
| Not Provided/Missing | 1 | 0 |
| Race |  |  |
| Caucasian | 31 | 30 |
| African American | 3 | 7 |
| Asian American | 2 | 1 |
| Bi/Multiracial | 2 | 4 |
| Not Provided/Missing | 2 | 0 |
| Average Annual Income |  |  |
| < \$5,000-\$19,999 | 10 | 11 |
| \$20,000-\$34,9999 | 9 | 7 |
| \$35,000 - \$59,999 | 8 | 5 |
| \$60,000 - \$99,999 | 5 | 6 |
| \$100,000 + | 3 | 10 |
| Not Provided/Missing | 5 | 3 |
| Sexuality |  |  |
| Heterosexual | 28 | 40 |
| Gay/Lesbian | 1 | 1 |
| Bisexual | 8 | 0 |
| Other | 1 | 1 |
| Not Provided/Missing | 2 | 0 |

*Note.* Samples were sex- and age-matched. PAI-BOR: Personality Assessment Inventory – Borderline subscale.

### Procedure

Participants underwent two semi-structured diagnostic interviews to assess for psychopathology and personality disorder symptoms (50,51). Interviews were administered by two trained research assistants who were supervised by the senior author. Participants in the BPD group met diagnostic criteria for three or more of the DSM-IV-TR BPD symptoms, an empirically derived threshold for identifying clinically significant symptoms (52). Exclusionary criteria for both groups included having a first-degree relative diagnosed with Bipolar I disorder or any psychotic disorder and a history of serious head injury or neurological disease. Control participants additionally had no history of psychiatric or substance abuse disorders.

In a separate session preceding the RS-fMRI scan and at six- and twelve-month follow-up visits, participants completed a battery of self-report questionnaires. We focuses here on the UPPS-P impulsivity scale (11,53) given the relevance of heightened impulsivity in adolescent BPD. The UPPS-P subscales measure (Positive and Negative) Urgency, (Lack of) Premeditation, (Lack of) Perseverance, and Sensation Seeking. Internal consistency was good to excellent at baseline and over follow-up (α_total_ = 0.95, α_mean-subscales_ = 0.88). We were particularly interested in the NU and PU subscales of the UPPS, given their relevance in psychopathology (13), though we compared our results with all UPPS scales to test specificity (Supplemental Methods and Results). All study procedures were approved by the Institutional Review Boards of the University of Pittsburgh (PRO13010486).

#### MR data acquisition

Data were acquired using a Siemens 3T Tim Trio scanner with a 32-channel head coil at the University of Pittsburgh Medical Center. We collected five minutes of resting-state fMRI data at the end of a broader scanning protocol; subjects were asked to keep their eyes open and relax, but not fall asleep. We used a simultaneous multi-slice echo-planar sequence sensitive to BOLD contrast with scanning parameters: TR = 1.0s, TE = 30ms, flip angle = 55°, voxel size = 2.3mm isotropic, 5x multiband acceleration. Participants completed a self-report questionnaire at the end of the protocol to determine if they fell asleep during the scan. No subjects were excluded for sleepiness.

### RS-fMRI preprocessing

RS-fMRI preprocessing was conducted within FSL, NiPy, and AFNI (54–56). Structural scans were registered to the MNI152 template (57) using affine and nonlinear transformations conducted in FSL. Functional image preprocessing included simultaneous 4-D motion and slicetiming correction (58), brain extraction, alignment of subject’s functional images to their anatomical scan using a boundary-based registration algorithm (59), and a one-step nonlinear warp to MNI152 space that concatenated functional-to structural, structural-to-MNI152, and fieldmap unwarping transformations. To mitigate motion-related artifacts we used ICA-AROMA (60), a data-driven classification algorithm that identifies and removes spatiotemporal components likely to reflect head movement. RS-fMRI data was not spatially smoothed for analysis (see Supplemental Methods) (61).

### Analytic approach

#### Nodal parcellation and functional connectivity matrix generation

To define our regions of interest, we parceled voxels into functional regions (nodes) by combining leading cortical and subcortical parcellations (Supplemental Methods) (62,63). We selected 19 fronto-striatal-limbic nodes, including portions of mPFC, OFC, and ACC as prefrontal nodes, and bilateral BLA, CeN, and VS as limbic nodes (Figure 1, Table 2). Prior to computing connectivity between nodes, we averaged the time series for voxels with reliable signal in each node to obtain a single nodal time series. For each subject and node, we prewhitened time series with an Auto-Regressive Moving Average (4,2) model, retaining the residual time series for FC estimation (Supplemental Methods).

**Figure 1.**
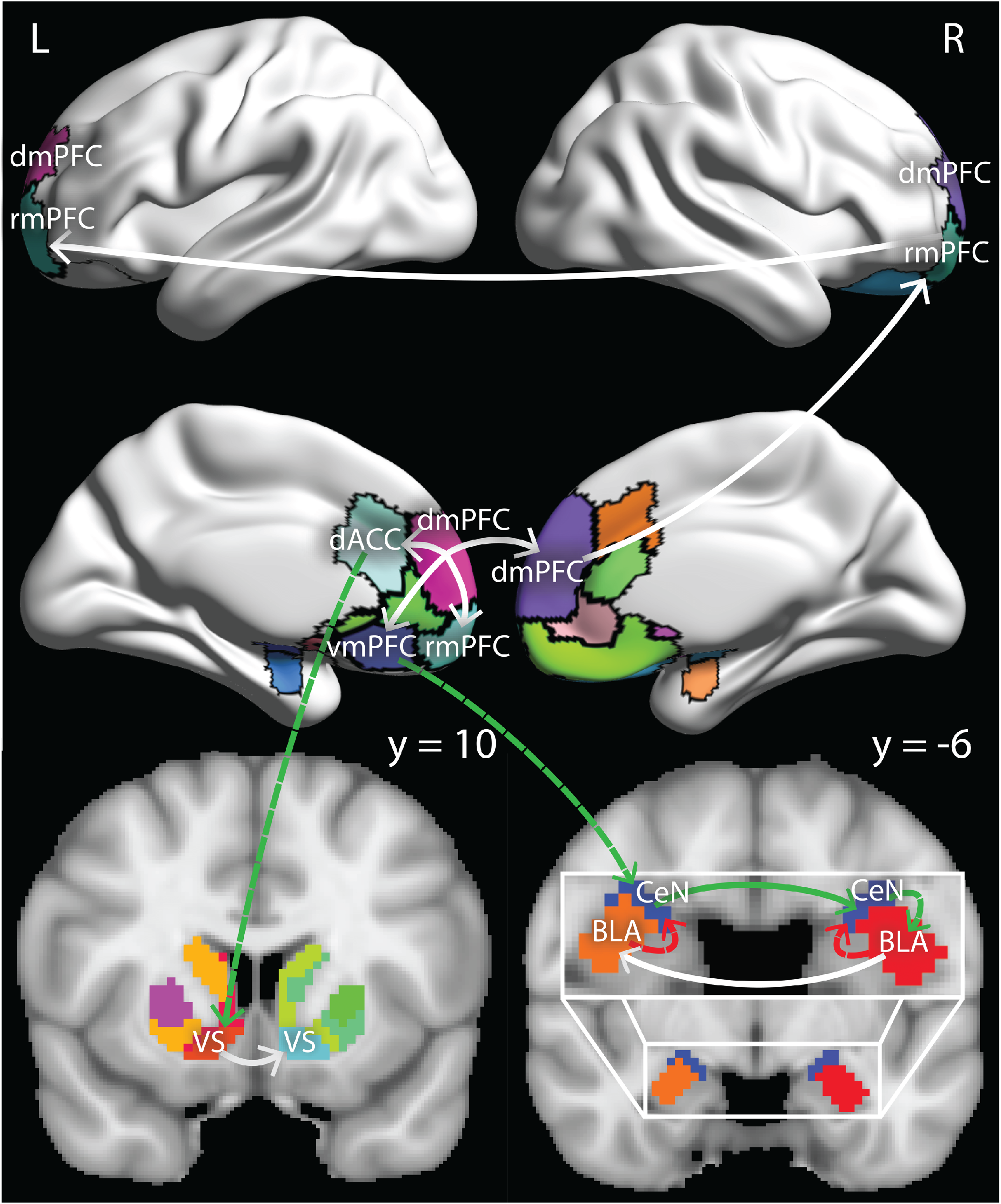
Graphical depiction of CS-GIMME results in anatomical space. *Note.* Nodes without labels were not fit with CS-GIMME but depict medial prefrontal and ACC nodes that were dropped from our initial consideration set. Arrows reflect the directed influence of one node on another at the group and subgroup (BPD vs control) level. Solid lines denote edges that were estimated at the *group (entire sample) level* with the solid green edge denoting a significantly higher edge values amongst subjects in the control group. Dashed lines denote edges estimated at the *subgroup (BPD vs HC) level,* with red and green dashed lines denoting edges that were only estimated for subjects in the BPD and control group, respectively.

**Table 2.** MNI center of mass coordinates for selected nodes

| Anatomical label | Parcellation label | CS-GIMME label | x | y | z |
| --- | --- | --- | --- | --- | --- |
| L OFC | 56 |  | -10 | 33 | -20 |
| L vmPFC | 84 | 1 | -6 | 35 | 10 |
| L rmPFC | 86 | 2 | -13 | 61 | -4 |
| L rACC | 88 |  | -6 | 43 | 8 |
| L dmPFC | 89 | 3 | -9 | 57 | 20 |
| L dACC | 90 | 4 | -6 | 29 | 25 |
| R OFC | 159 |  | 12 | 37 | -21 |
| R rmPFC | 161 | 5 | 15 | 63 | -6 |
| R dACC | 180 |  | 7 | 29 | 29 |
| R vmPFC | 191 |  | 5 | 36 | -14 |
| R rACC | 192 |  | 8 | 41 | 5 |
| R dACC | 193 |  | 5 | 28 | 16 |
| R dmPFC | 194 | 6 | 9 | 56 | 19 |
| L BLA | 201 | 7 | -25 | -5 | -22 |
| R BLA | 202 | 8 | 25 | -3 | -22 |
| L CeN | 203 | 9 | -20 | -6 | -15 |
| R CeN | 204 | 10 | 19 | -5 | -15 |
| L VS | 215 | 11 | -13 | 12 | -8 |
| R VS | 220 | 12 | 11 | 13 | -8 |

*Note.* Includes MNI coordinates for the original 19-node set. Nodes with no CS-GIMME label were excluded from EC graph estimation based on non-significant FC differences between groups.

Finally, we computed Pearson correlations among the pre-whitened time series to yield a 19 × 19 adjacency matrix for each subject representing undirected functional connectivity amongst fronto-limbic regions. In order to remove unreliable edges from these matrices, we applied a minimal consensus thresholding procedure (64). Specifically, we removed edges from all subjects that did not have a weight of *r* = .1 or higher in 25% or more of subjects. This resulted in the removal of 17 edges (10%) that were concentrated in OFC-subcortical and subcortical-subcortical connections.

#### Node selection: undirected analysis

The primary goal of our analyses was to examine EC using the Confirmatory Subgrouping Group Iterative Model Multiple Estimation algorithm (CS-GIMME) (65,66). However, given that GIMME conjointly estimates conditional relationships among all nodes, the number of free parameters increases exponentially as the number of nodes increases and parameter reliability decreases (67). To promote model convergence and reliable EC estimation, we performed a node selection analysis by fitting a single logistic ridge regression model predicting group status by all undirected edges, retaining nodes with edges that jointly predicted group status at the *p* < .01 level and were thus most likely to be implicated in EC (Supplementary Methods and Results) (68,69).

#### Effective connectivity network estimation and relations to group status and age

We retained the preprocessed time series of twelve^2^ fronto-limbic nodes based on our node selection analysis (Table 2; Supplemental Methods and Results). We estimated EC between these nodes using the CS-GIMME algorithm, a recent extension of the GIMME algorithm that reliably detects the presence and direction of edges in fMRI data at the individual, group, and sample levels.

After obtaining directed graphs from CS-GIMME, we investigated group differences in the role of individual nodes (nodal centrality) in fronto-limbic circuits. We calculated in- and out-degree centrality for each node that showed evidence of incoming or outgoing edges in the bestfitting CS-GIMME model (denoting the summed score of incoming and outgoing edges for each node; Supplemental Methods). In order to identify which nodal centrality estimates best differentiated groups, we entered these into a single logistic regression^3^ predicting group status:

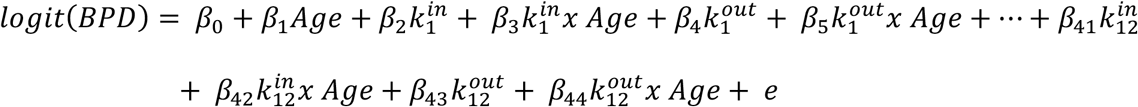

where 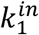 is in-degree for node 1 (Table 2).

#### Predicting stability and change in impulsivity

We tested for group differences in baseline ERI in addition to within-person changes in ERI symptoms over 6- and 12-month follow-up. We first fit a latent growth curve model (LGCM) (71), modeling latent intercept and slope terms for both NU and PU^4^, which describe the baseline level and rate of within-person change, respectively, in NU and PU scales. In this model, group, age, and their interaction were included as predictors of LGCM variables (Fig 2).

**Figure 2.**
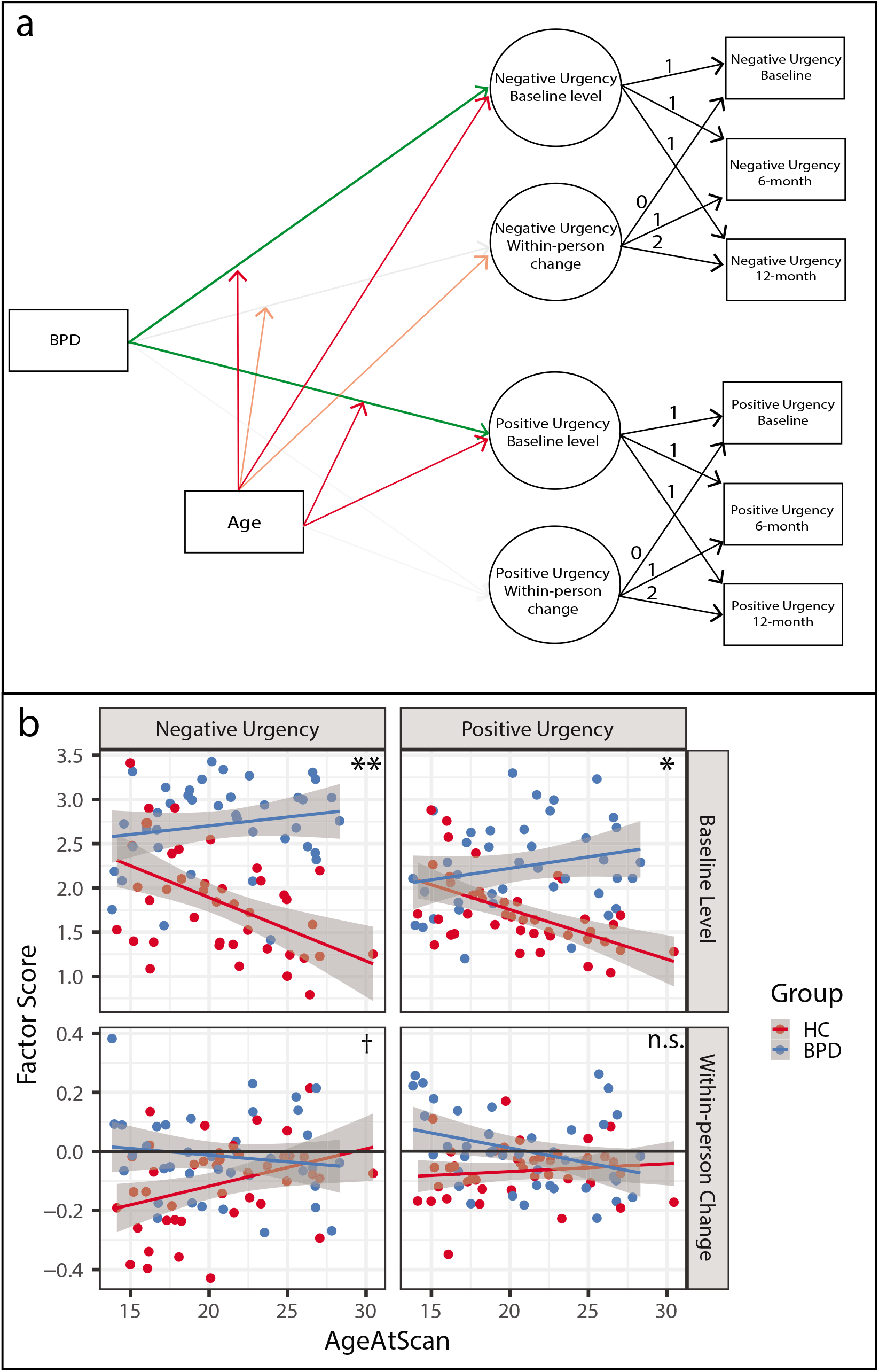
Latent growth curve model and relations to group and age. *Note.* a) Conceptual diagram of latent growth curve model reported in Table 5. Solid green lines denote positive parameter estimates, demonstrating that group main effects are higher for both intercept terms (overall levels of ERI) in the BPD group. Solid red lines denote negative parameter estimates, demonstrating that age main effects on both intercept terms indicate decreasing levels of ERI in the total sample. Faded red lines denote “marginally” significant parameter estimates (0.05 < *p* < 0.10). Paths from age that intersect with paths from BPD to LGCM terms represent interactive effects (moderation) of age and group to predict LGCM terms. b) Visual depiction of age x group interaction for each LGCM term, demonstrating significant group x age interactions for both NU and PU intercept terms and a marginally significant group x age interaction for NU slope.

We then tested if group and group-by-age differences in the longitudinal model of ERI were mediated by EC estimates of the R CeN (Fig 3). As detailed below, the results from the joint logistic model indicated that R CeN played a particularly important role in differentiating BPD from HC participants (Table 4). Thus, we fit a combined dual-mediator LGCM, which tested the hypothesis that directed functional input and/or output of the R CeN mediated the relationship between group status and latent intercept and slope variables. In our final combined model, both in- and out-degree of the R CeN were mediators of LGCM variables for both NU and PU. We allowed age to predict levels of directed centrality of R CeN, and to moderate the relationship between centrality estimates and LGCM variables (73).

**Figure 3.**
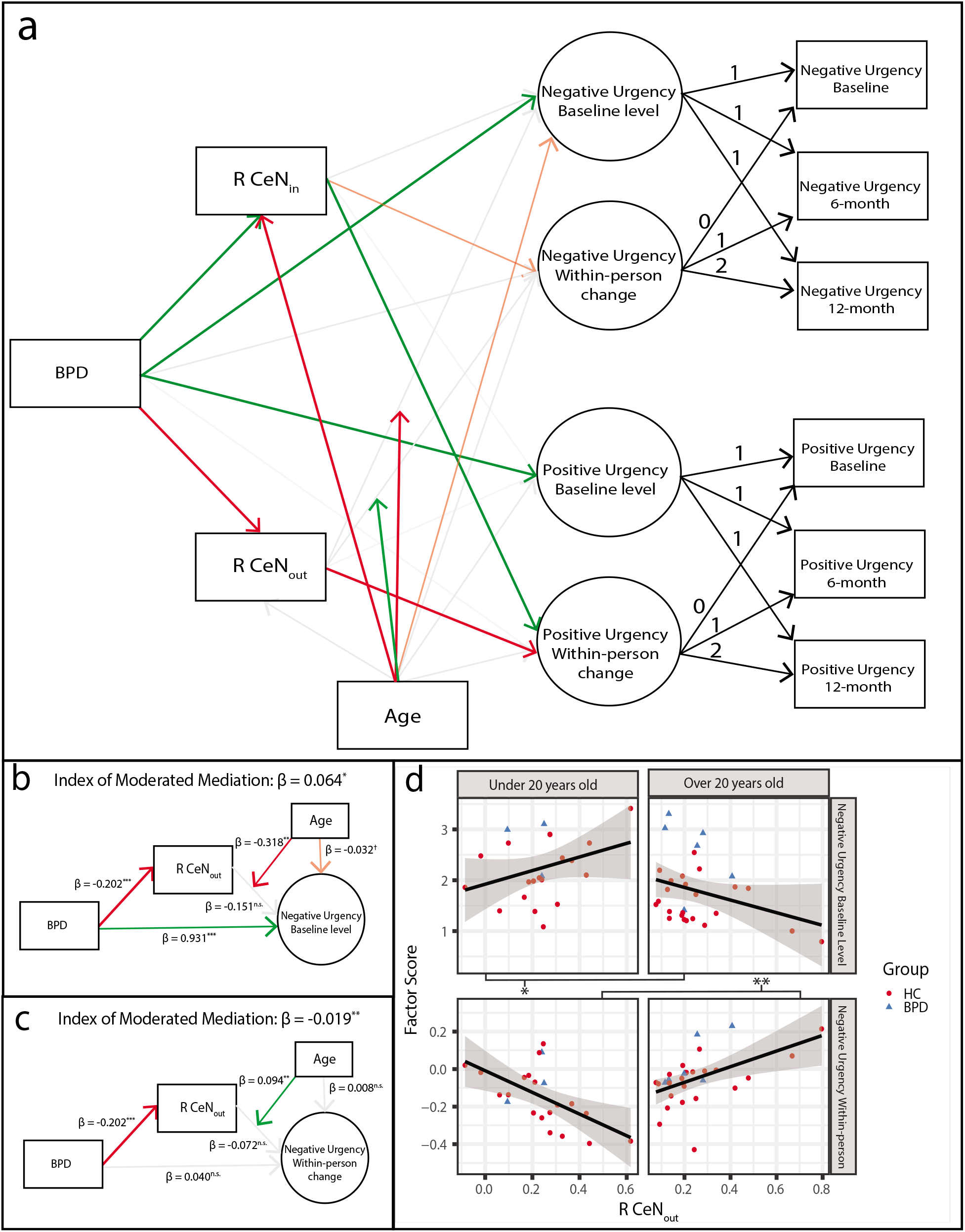
latent growth curve model with dual mediators reveals group differences in stability and change in ERI is underpin by intrinsic directed connectivity of R CeN. *Note.* a) Conceptual diagram of final latent growth curve model reported in Table 6. Solid green lines denote positive parameter estimates, demonstrating that main effects are higher for both baseline ERI variables and R CeNin in the BPD group. Solid red lines denote negative parameter estimates, demonstrating that R CeNin decreased with age, and that BPD was associated with lower R CeNout, which was in turn associated with lower (e.g. “more negative”) within-person change in PU . Faded red lines denote “marginally” significant parameter estimates (0.05 < *p* < 0.10). Paths from age that intersect with paths from R CeNin and R CeNout to LGCM terms represent interactive effects (moderation) of age on the second-stage mediation (i.e. from R CeN connectivity to LGCM terms). b) Significant moderated mediation of from BPD to NU baseline levels through R CeNout, which was moderated by age. c) Significant moderated mediation of from BPD to NU within-person change through R CeNout, which was moderated by age. d) Visual split of moderated mediation into young participants (under 20 years old) and older participants (over 20 years old). Note most importantly the lower left panel, showing that amongst adolescents, participants with higher R CeNout had more negative levels of within-person change in NU over one year, demonstrating more rapid reductions in NU symptoms over the one-year follow-up.

**Table 3.** Directed edge values from CS-GIMME

| Node from | Node to | Edge value (SD):<br>Whole Sample | Edge value (SD):<br>BPD | Edge value (SD):<br>Control | *p* |
| --- | --- | --- | --- | --- | --- |
| L dmPFC | L vmPFC | 0.461 (0.21) | 0.445 (0.23) | 0.477 (0.20) | .49 |
|  | L rmPFC | 0.437 (0.24) | 0.458 (0.18) | 0.398 (0.29) | .26 |
|  | L dACC | 0.534 (0.21) | 0.517 (0.18) | 0.550 (0.24) | .49 |
|  | R dmPFC | 0.818 (0.14) | 0.841 (0.12) | 0.798 (0.15) | .15 |
| R rmPFC | L rmPFC | 0.165 (0.15) | 0.186 (0.16) | 0.145 (0.14) | .22 |
| R dmPFC | R rmPFC | 0.380 (0.26) | 0.412 (0.28) | 0.350 (0.24) | .29 |
| R BLA | L BLA | 0.232 (0.14) | 0.220 (0.15) | 0.244 (0.13) | .44 |
| L CeN | R CeN | 0.200 (0.13) | 0.169 (0.14) | 0.228 (0.13) | .05* |
| L VS | R VS | 0.273 (0.13) | 0.258 (0.13) | 0.289 (0.12) | .27 |
| L vmPFC | L CeN |  |  | 0.127 (0.09) |  |
| L dACC | L VS |  |  | 0.211 (.14) |  |
| R CeN | R BLA |  |  | 0.208 (.13) |  |
| L BLA | L CeN |  | 0.153 (.13) |  |  |
| R BLA | R CeN |  | 0.163 (.12) |  |  |

*Note.* Mean edge values of directed edges estimated by CS-GIMME, calculated on the whole sample and split by group. P-values are based on t-tests between groups. The lower section includes subgroup-specific edges.

**Table 4.** Joint model: group differences in in- and -out degree centrality

| Node (CS-GIMME label) | Centrality | Est.(S.E.) | t-score (p value) |
| --- | --- | --- | --- |
| L vmPFC (1) | In | -0.097(.060) | -1.62 (.11) |
|  | Out | -0.025(.053) | -0.481 (.63) |
| L rmPFC (2) | In | -0.044(.078) | -0.56 (.57) |
|  | Out |  |  |
| L dmPFC (3) | In |  |  |
|  | Out | 0.141(.110) | 1.28 (.21) |
| L dACC (4) | In | -0.029(.062) | -0.46 (.65) |
|  | Out | -0.124(.064) | -1.94 (.06)<sup>†</sup> |
| R rmPFC (5) | In | 0.028(.083) | 0.34 (.74) |
|  | Out | -0.029(.055) | -0.54 (0.59) |
| R dmPFC (6) | In | -0.043(.053) | -0.82 (.42) |
|  | Out | -0.008(.074) | -0.11 (.91) |
| L BLA (7) | In | -0.006(.061) | -0.10 (.92) |
|  | Out | 0.026(.064) | 0.37 (.71) |
| R BLA (8) | In | 0.060(.060) | 1.00 (.32) |
|  | Out | 0.089(.076) | 1.18 (.24) |
| L CeN (9) | In | 0.039(.072) | 0.54 (.59) |
|  | Out | -0.082(.060) | -1.35 (.18) |
| **R CeN (10)** | **In** | **0.149(.061)** | **2.47 (.02)<sup>\*</sup>** |
|  | **Out** | **-0.217(.057)** | **-3.82 (< .001)<sup>\*\*\*</sup>** |
| L VS (11) | In | -0.055(.064) | -0.87 (.39) |
|  | Out | -0.032(.065) | -0.49 (.62) |
| R VS (12) | In | -0.010(.062) | -0.17 (.87) |
|  | Out |  |  |

*Note.* Rows with missing values indicate that there was no group- or subgroup-level edge entering/exiting the node, and thus we elected to not compute estimates of centrality based only on individual-level edges. Results represent coefficients in a single model that were allowed to compete to explain variation in group status. Note that positive coefficients represent that degree centrality estimates were higher in the BPD group. We also ran a supplementary set of analyses that fit separate models to each effect (Supplemental Methods and Results, Table S2), which provides an intuition of the marginal group effects, whereas the joint model account the complex correlational structure of these connectivity scores (Fig S3).

## Results

### Directed network estimation

A graphical depiction of the best-fitting CS-GIMME network is provided in Figure 1; directed edge statistics are listed in Table 3. Effective connectivity estimates from CS-GIMME generated positive directed edges at both the group level (nine total) and the subgroup level (five total, two in the BPD group). We found a range of directed edges from the dmPFC to several mPFC nodes at the group level including left dACC, rmPFC, and vmPFC, as well as right dmPFC. We also found evidence at the group-level for EC between bilateral CeN, VS, and BLA nodes. In healthy controls, but not BPD participants, CS-GIMME detected edges from dACC to VS, vmPFC to the CeN, and CeN to BLA. Conversely in BPD subjects, the CS-GIMME algorithm found BPD-specific edges from bilateral BLA to CeN.

### Group and age-related differences in fronto-limbic connectivity

In a joint model predicting group status from directed centrality estimates we found a dissociation between in- and out-degree centrality in R CeN by group (Table 4). Subjects in the BPD group had significantly higher in-degree (*t* = 2.47, *p* = .02) and significantly lower out-degree (*t* = −3.82,*p* < .001) of R CeN. This effect did not differ by age: all age-related terms in the joint model were nonsignificant.

### Associations with ERI scales

As expected, NU and PU subscale scores were significantly higher in the BPD group at baseline compared to controls (*β*_NU_ = 0.88, *p* < .001 *β*_PU_ = 0.53, *p* < .001; Table 5, Fig 2). We also found group x age interactions in baseline ERI scores: older controls had lower baseline NU and PU scores than younger controls, whereas baseline NU and PU scores in the BPD group did not differ substantially by age (*β*_NU_ = 0.10, *p* = .01; *β*_PU_ = 0.086, *p* = .01).

**Table 5.**
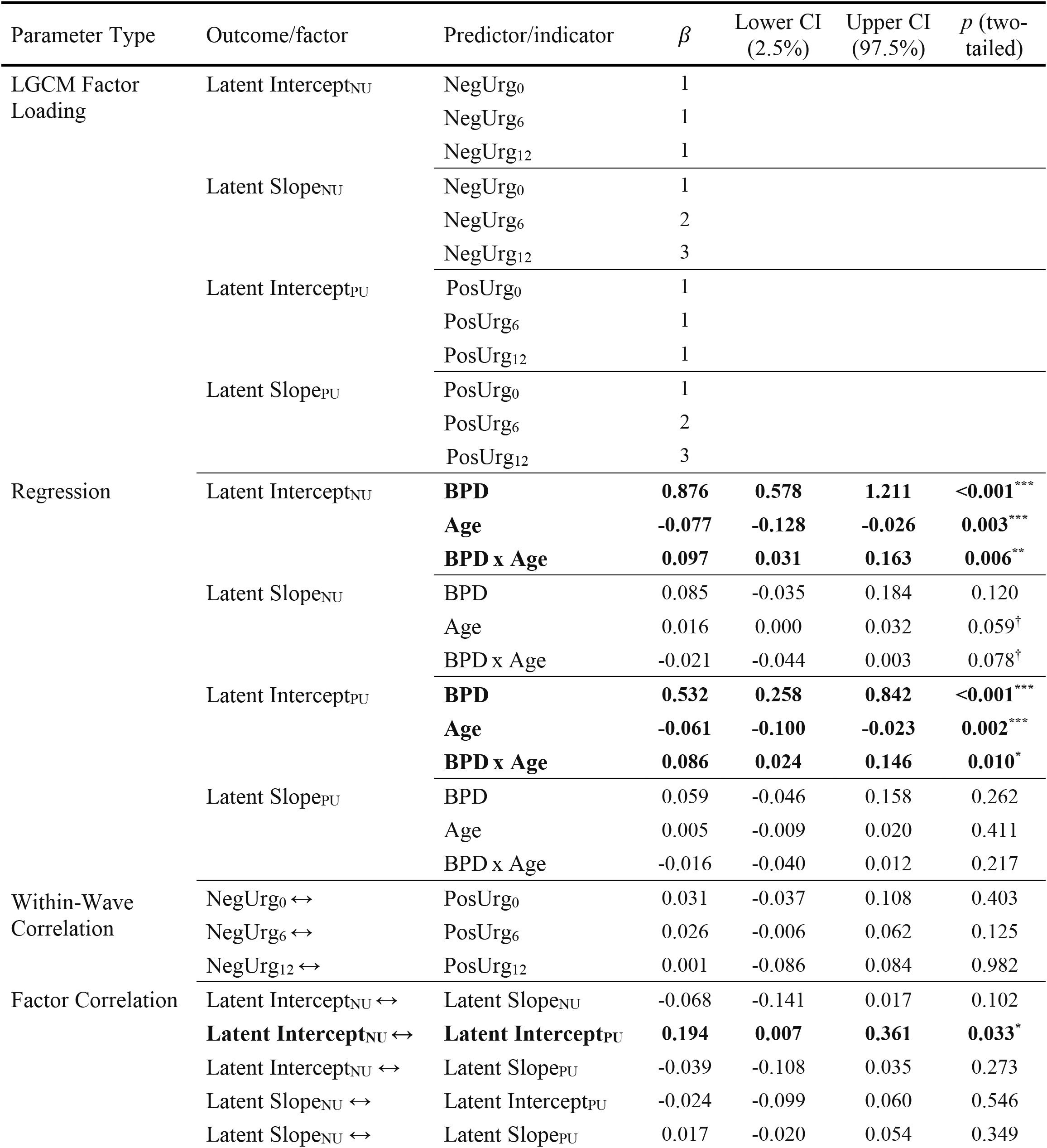
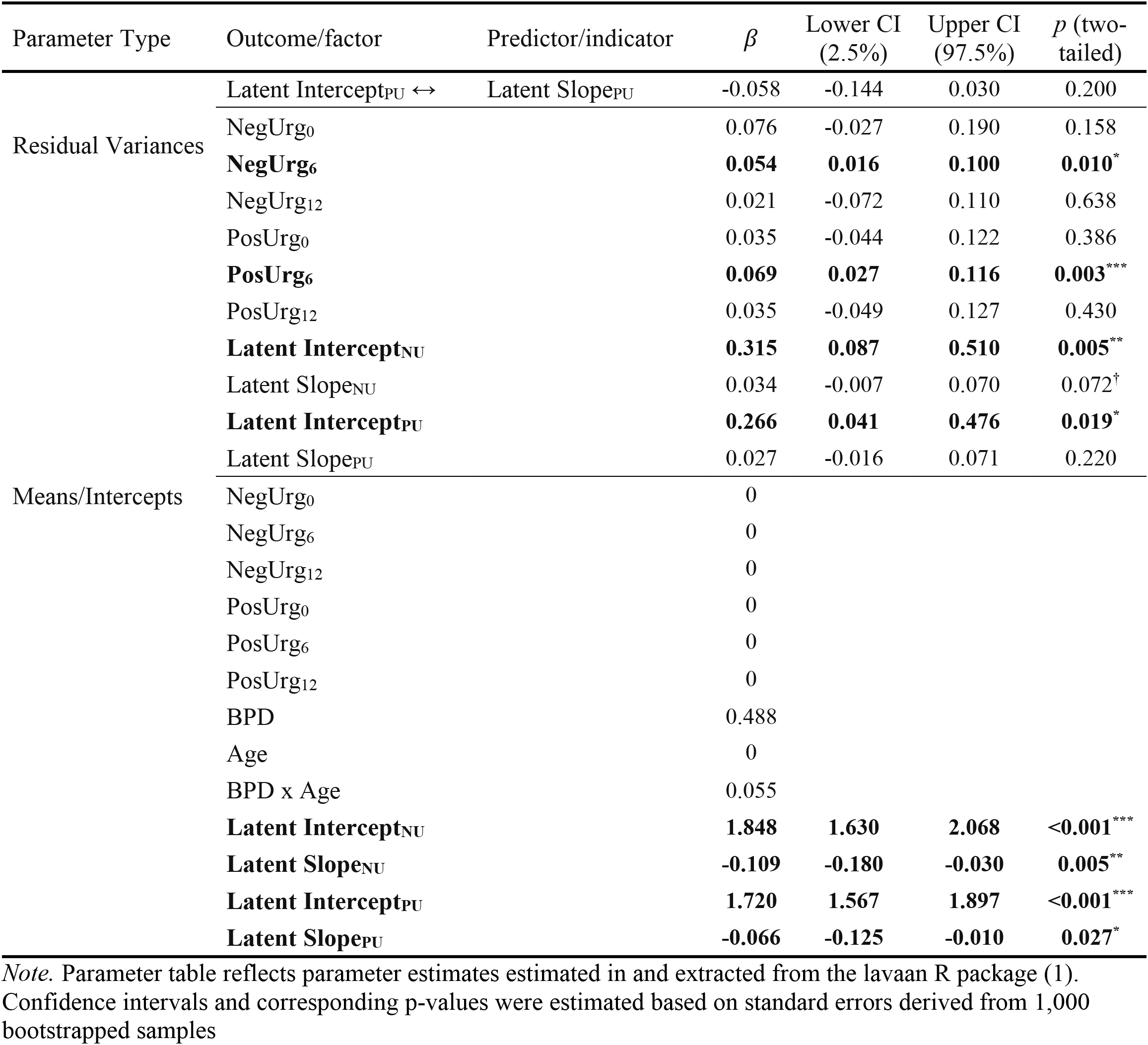
Parameter table: latent growth curve model of ERI scales and relations to BPD, Age, and their interaction

In a final combined LGCM, we tested if group differences in ERI scores over the one-year follow-up period were accounted for by effective connectivity of R CeN at baseline. Results are detailed in Table 6 and visually depicted in Fig 3. R CeN out-degree statistically mediated the association of group status with within-person changes in PU (Indirect Effect = 0.08; *p* = 0.04), such that CeN out-degree was lower in the BPD group *(β* = −0.20 p < .001) and was related to sharper within-person declines in PU *(β* = −0.35, p = .03). For NU, we found that the influence of R CeN out-degree on baseline level and within-person change in NU depended on age: Hayes’ index = 0.06 *(p* = .01) and −0.02 *(p* = .01), respectively. In younger participants, higher R CeN out-degree was associated with higher baseline NU and greater within-person decreases in NU over the one-year follow-up whereas in older participants the opposite pattern was observed (Fig 4b-d).

**Table 6.**
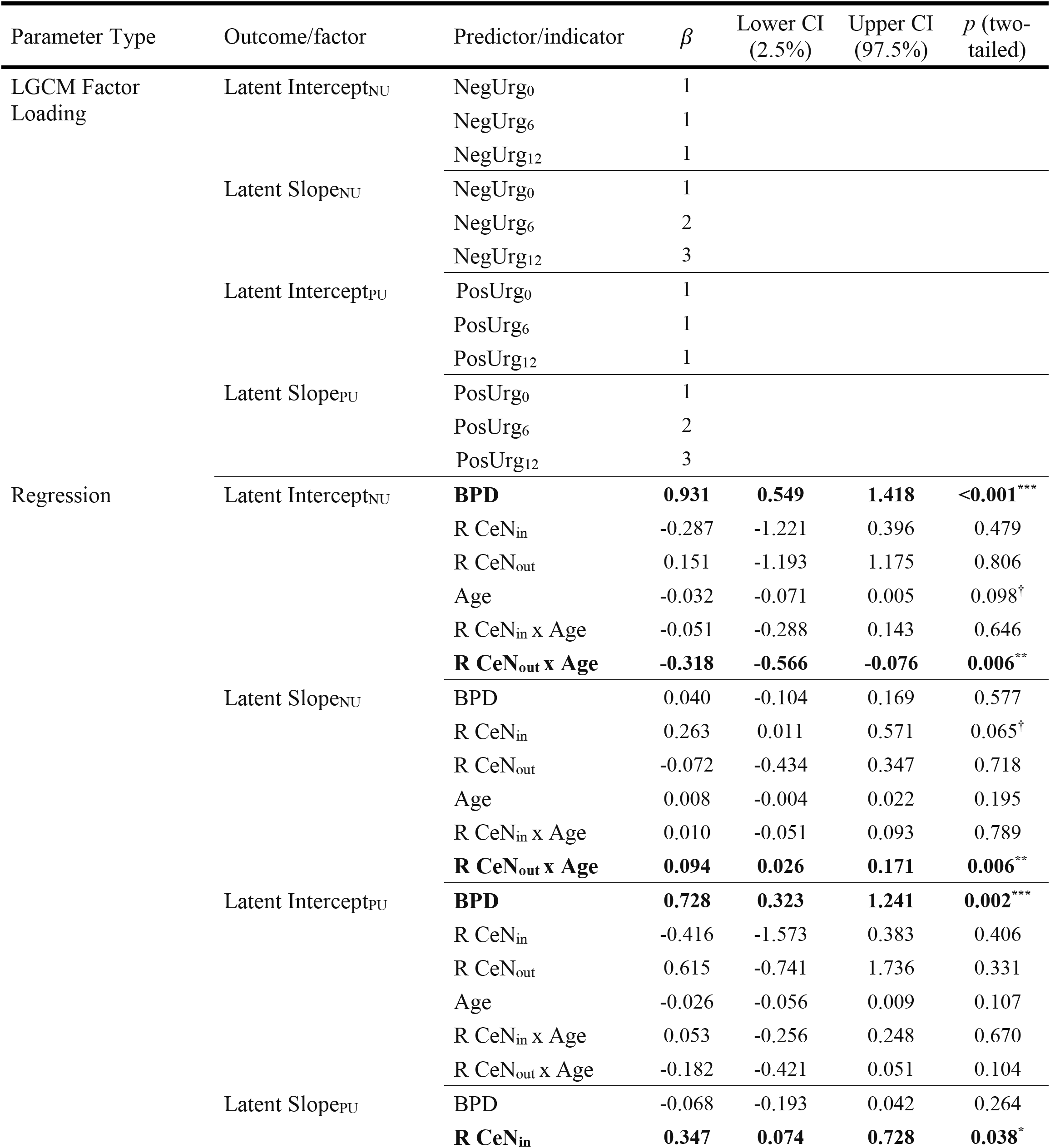
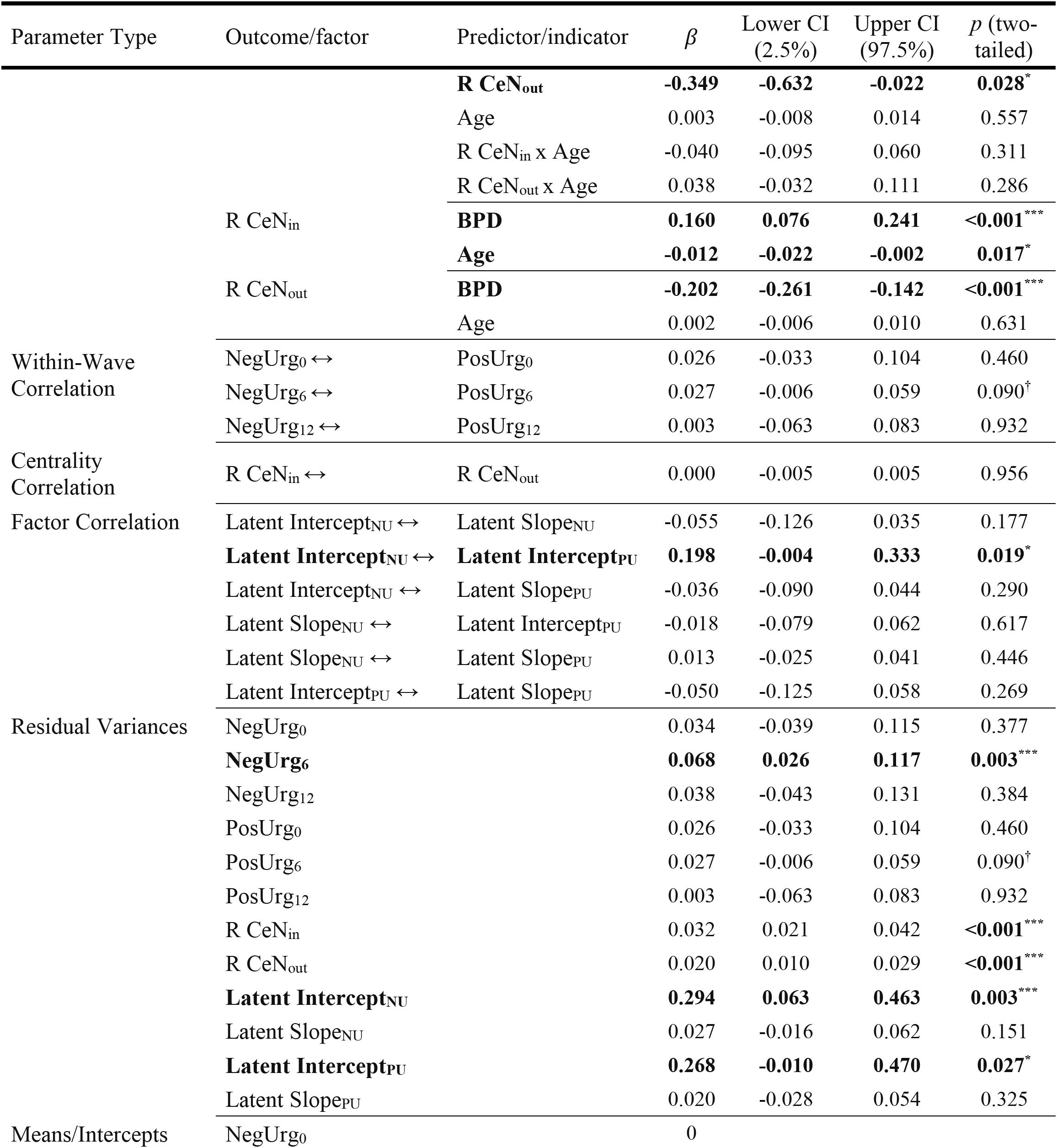
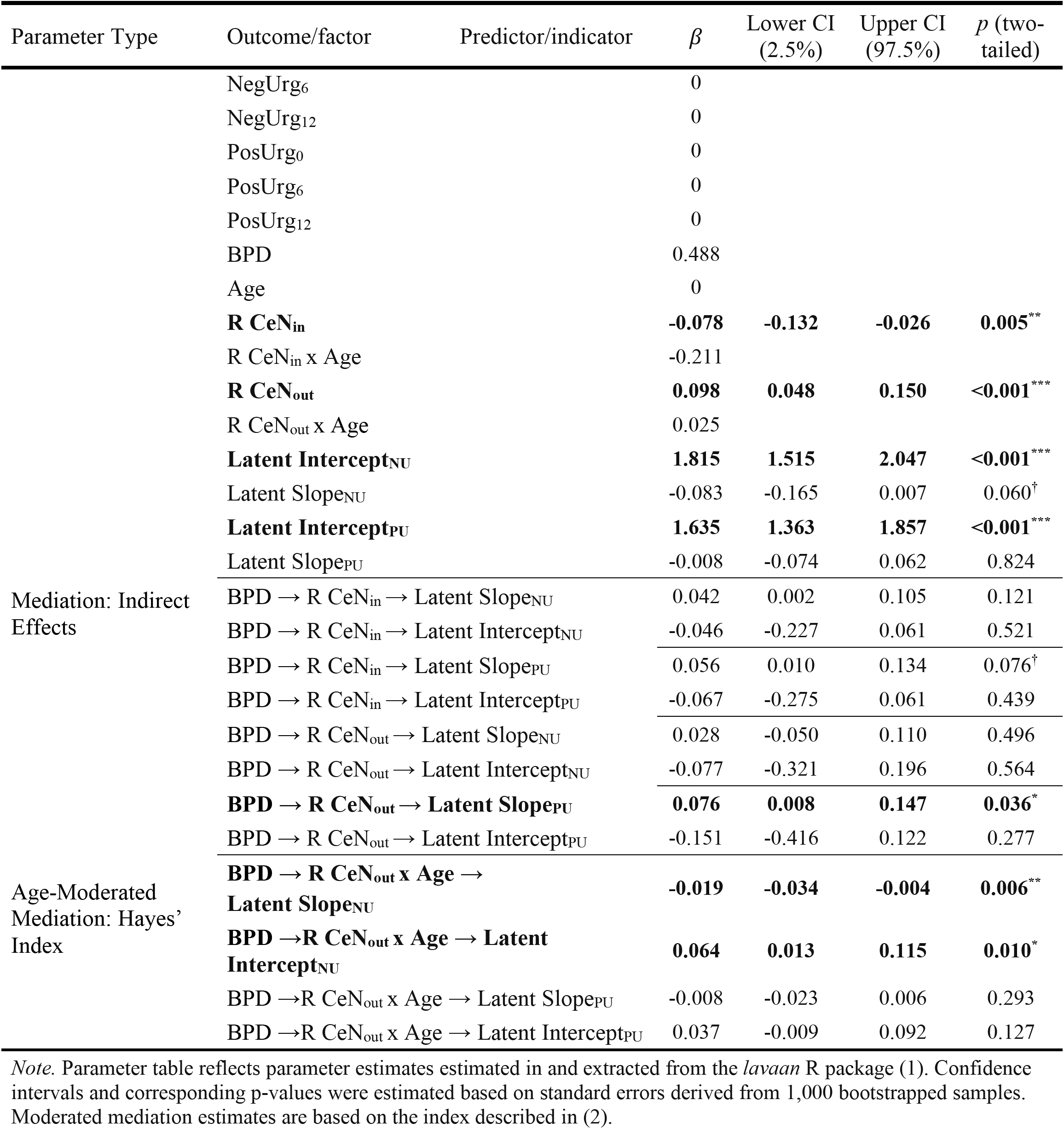
Parameter table: dual-mediator latent growth curve model includes mediation by R CeN in- and out-degree

Although our emphasis on ERI was informed by clinical theory and prior BPD research, we also note that in preliminary mixed-effects models, we found significant group by age interactions in levels of NU and PU but not other facets of impulsivity (Supplemental Methods and Results; Table S3).

## Discussion

In a sample of adolescents and emerging adults with BPD symptoms and matched healthy controls, we found that resting-state effective connectivity (EC) of amygdala subnuclei at study baseline accounted for group differences in initial levels and within-person change in ERI over a one-year follow-up period. Specifically, input to R CeN (in-degree) was significantly higher in the BPD group, which largely reflected a heightened influence of BLA on CeN. Conversely, in the control group, higher levels of output of R CeN (out-degree) were primarily attributable to R CeN’s directed input to BLA (Figs S4-5). Crucially, output of R CeN statistically mediated the association between BPD symptoms and baseline ERI, as well as within-person stability of ERI in borderline adolescents. Our results indicate that CeN plays an important role in impulsive behaviors in response to intense emotions. Importantly, whereas we found evidence of directed connectivity from vmPFC to CeN in controls, this functional connection was not reliably observed in BPD participants. This suggests an altered integration of cortico-limbic signals in the CeN, which has important clinical implications regarding the developmental course of BPD from adolescence to emerging adulthood.

Fronto-limbic disturbances are frequently reported in fMRI studies of adults with BPD, with the strongest evidence for amygdala hyperactivity in the processing of emotions. Further, prior evidence suggests that the functional interaction between vmPFC and amygdala is suppressed in individuals with BPD when presented with emotional stimuli (20,23). We found that EC from vmPFC to CeN was *positive* in controls, but absent in the borderline group. This positive connection in our data is in line with other BOLD EC studies showing positive EC from vmPFC to amygdala (78–80), though we note that animal studies of fear conditioning find that mPFC/infralimbic projections to the amygdala are primarily inhibitory (74–77). One possibility is that positive EC between these regions reflects an increased *capacity* of vmPFC to control affective responding in CeN (79). This interpretation aligns well with evidence that humans with vmPFC lesions show potentiated amygdala responsivity (81). Additionally, preclinical research indicates that electrical stimulation of mPFC neurons suppress the activity of output neurons in CeN that control autonomic/emotional arousal (75). If functional connections between vmPFC and CeN are absent or significantly weakened in young people with BPD, excitatory connections from BLA to CeN (30,82) could exert heightened influence over CeN efferents that control arousal.

While CeN encodes the general affective and motivational significance of emotional events (83,84), BLA is involved in assigning emotional significance to sensory stimuli (85,86). BLA projections to CeN control the expression of anxious behavior (82), consistent with the interpretation that heightened BLA-to-CeN EC in BPD adolescents could reflect a stronger tendency to initiate impulsive behaviors in response to emotional arousal. A heightened ability of BLA to influence CeN in BPD may reflect a tendency to translate sensory signals with greater emotional significance (BLA) (87,88) to a state of enhanced physiological arousal (CeN). Such high arousal states could predispose adolescents towards making impulsive choices, either to enhance positive emotion or to escape negative emotion.

Crucially, differential EC between the R CeN and BLA was associated with self-reported ERI levels at baseline and within-person change in ERI over 12-month follow-up (Fig 3). Adolescence is associated with increased levels of impulsivity, which decrease in the general population, yet adolescents in the BPD group demonstrated greater stability in ERI symptoms. Within-person decreases in NU in healthy adolescents (Fig 3b, lower left) were associated with stronger EC from R CeN to R BLA at baseline. Most studies find that functional interactions between BLA and CeN reflect the influence of BLA activity on CeN, rather than the other way around (27,82). Thus, our finding that R CeN output mediates within-person decreases in NU may reflect an increased capacity to resist BLA modulation in controls (a capacity that may be supported by signaling from vmPFC). Given that EC from CeN to BLA was not observed in the BPD group, we propose that BLA control of the CeN is one candidate mechanism for explaining the stability of ERI in BPD. This proposal extends earlier findings of fronto-limbic abnormalities in BPD by illustrating a more anatomically nuanced account of intra-amygdala information flow than previously described in this population. Most importantly, we demonstrate that intraamygdala EC in our data predicted the clinical course of ERI symptoms up to one-year postscan.

To our knowledge, this is the first study in humans to demonstrate tradeoffs in EC between BLA and CeN in any psychiatric population. Furthermore, we leveraged recent developments in effective connectivity estimation (66) that are well-suited for detecting group-specific connectivity patterns. Our findings have clear implications for future study in other disorders with heightened levels of ERI including addiction (14), anxiety disorders (89), eating disorders (90), and PTSD (91). As such, it is worth noting that dissociations in functional (undirected) connectivity of BLA and CeN have been previously documented in human subjects with these disorders (14,92–94).

A few limitations are worth noting. First, our RS-fMRI acquisition was cross-sectional. Though we demonstrated the ability of a cross-sectional measure of brain connectivity to explain within-person changes in ERI over 12 months, longitudinal neuroimaging studies offer the opportunity to study developmental changes in EC between BLA and CeN. Second, while relating self-reported ERI levels to measures of intrinsic EC is an important descriptive step in identifying candidate neural circuits implicated in personality pathology, highly stable levels of ERI in the BPD group may be due to selection criteria for our study. That is, ERI stability in the BPD group may reflect a byproduct of selecting participants with heightened BPD symptoms (including impulsivity). Finally, future studies should include clinical comparison groups to clarify this specificity/generality of our findings, as differences in fronto-limbic connectivity have been documented across many disorders.

We present evidence that effective connectivity between vmPFC, CeN, and BLA is altered in adolescents and emerging adults with BPD symptoms. One speculative interpretation is that mPFC and BLA compete in modulating CeN activity, and differential contributions of these regions to CeN activity underlie stability and change in ERI symptoms from adolescence through emerging adulthood. Further, our findings demonstrate altered fronto-limbic connectivity in adolescents with BPD symptoms, including a functional disconnection between vmPFC and CeN and tradeoffs in control between functionally distinct subnuclei of the amygdala, which may underlie impulsive behaviors in the face of strong emotions. We hope that future studies build on our results by examining biomarkers that can inform treatments for adolescents at high risk for negative BPD-related outcomes in adulthood.

## Acknowledgments

This work was funded by the National Institutes of Mental Health (K01 MH097091 and_R0l MH119399 to MNH). The funding agency had no role in the design and conduct of the study; the collection, management, analysis, and interpretation of the data; the preparation, review, and approval of the manuscript; or the decision to submit the manuscript for publication.

## Footnotes

1 While no studies have *specifically* examined stability and change in ERI during the transition from adolescence to early adulthood, impulsivity in these studies is contrasted to sensation seeking, which shows clear differences in developmental course and relevance to psychopathology (45,46).

2 We noted previously, that GIMME may struggle with parameter identifiability with a large number of nodes (approximately 20 nodes) (70). To ensure the robustness of our results we re-estimated directed edges with CS-GIMME on the original a priori 19-node set. and retained nearly identical results, with the exception of a small number of intra-mPFC edges estimated with the inclusion of additional right mPFC nodes. Fronto-limbic connectivity was preserved across analyses, indicating that our choice to trim the number of nodes used in CS-GIMME estimation did not bias results.

3 In order to get a broad sense of group and age-related associations, preliminary analyses that were not jointly fit are described in Supplemental Methods and Results and summarized in Table S2. However, the joint approach is preferred as a straightforward correction for multiple comparisons and addresses the conditional associations amongst centrality metrics (Figure S2).

4 Simultaneously modeling the growth of two separate variables is considered an instantiation of a parallel process growth curve model (72), where correlations within measurement waves help to further reduce unexplained variation in the model. These models also allow for latent intercepts and slopes to predict one another, though in our analysis, we elected to leave growth parameters simply correlated with one another, as there is no strong evidence in the literature that would predict that levels of NU or PU to have a direct influence on the other.

