## Supplementary material for "Dissociation in basolateral and central amygdala effective connectivity predicts the stability of emotion-related impulsivity in adolescents with borderline personality symptoms: a resting-state fMRI study": RS_FL_BPS_supplement.pdf

### Supplemental Methods

#### fMRI QA and exclusion

For all participants, we calculated volume-to-volume framewise displacement (1). We excluded subjects with FD > 0.5mm in at least 20% of the volumes, or any FD > 10mm. This led to the removal of six participants, four of whom were in the BPD group. We removed one additional subject from the BPD group whose data did not pass serial residual correlation checks after pre-whitening our data. We further removed one participant from the BPD group who passed our head motion criteria but whose functional connectivity matrix was remarkably different from the group average (see Fig S1).

#### RS-fMRI preprocessing procedures

Although many resting-state fMRI studies apply spatial smoothing, we analyzed unsmoothed data because 1) our whole-brain parcellation encompasses the entire brain volume, leading to the close spatial proximity of many nodes, and 2) a recent report documenting that spatial smoothing alters graph theory-based measured of connectivity non-uniformly across the brain (2). Although our connectivity analyses were conducted on unsmoothed data, ICA-AROMA was conducted on data that were spatially smoothed with a 5mm FWHM gaussian kernel (FSL *susan*), consistent with recommended guidelines. Specifically, spatial smoothing increases SNR in BOLD data, allowing for an increased ability to detect structured artifacts that should be removed from the signal (such as components related to subject movement; 1). Based on the results of ICA-AROMA, we regressed motion-related components out of the unsmoothed data using *fsl regfilt* (i.e. “non-aggressive” denoising). AROMA’s automated component selection approach has recently been shown to be superior to other competing procedures in removing motion artefacts while preserving the signal of interest, and it largely eliminates distance-dependent motion-FC correlation effects (3–5).

**Analytic approach**

**Parcellation of brain regions into functional nodes** Our parcellation was based on the 200-node cortical parcellation derived recently by (6). The study authors used RS-fMRI data from 1489 participants and demonstrated that their parcellation provided a more homogenous partition of whole-brain neural activity compared with four leading cortical parcellations and agreed well with cortical boundaries. For amygdala nodes, we used the Harvard-Oxford subcortical atlas, retaining the bilateral CeN and BLA regions. Bilateral ventral striatum nodes were extracted from the anatomical mask provided by (7).

We chose to focus our analysis on medial prefrontal structures rather than the entire PFC to reflect prior knowledge of cortico-limbic connectivity. While documented FC differences between the dlPFC and amygdala have been observed in BPD (8), retrograde and anterograde tracing studies of amygdala-PFC connections support the presence of direct mPFC-amygdala projections (particularly in vmPFC), though not for dlPFC (9). As such we focused on mPFC and ACC in the current analysis to limit our FC estimates to edges between nodes that have a reasonable chance of having direct anatomical links, though this is impossible to directly verify solely based on the BOLD signal.

After creating our combined parcellation we calculated subject-level masks that reflected the proportion of voxels in each ROI that contained unreliable signal, as indicated by voxelwise standard deviation equal to zero and/or all values equal to zero. Visual inspection of the subject-level masks indicated that problematic voxels were located predominantly in inferior temporal regions and to a lesser extent, orbitofrontal regions, likely reflecting signal loss due to susceptibility artifacts. We then merged all binary masks into a group-level mask with voxel-wise values equal to the proportion of subjects with reliable signal in the voxel. In order to ensure that ROI time series reflected the same voxels across participants, we removed all voxels from the parcellation in which less than 95% of subjects had reliable signal. This procedure removed 600 voxels —approximately .7% of the total voxels — from our parcellation.

**Pre-whitening and adjacency matrix generation.**

After performing motion removal procedures and constructing our combined parcellation, we pre-whitened our nodal time series using a series of increasingly complex ARMA models (10) prior to calculating functional connectivity to ensure that cross-correlation estimates were not biased by the temporal (i.e. autocorrelated) structure of BOLD data (11,12). This decision was based on the concern that failing to remove autoregressive components of fMRI time-series violates a key assumption of the general linear model (specifically, GLM residuals must be *iid* and normally distributed) (1). Furthermore, estimates of cross-correlation can be misestimated when time series have similar autoregressive properties that do not reflect true interregional connectivity. To overcome this concern, auto-regressive models including ARMA have received increasing attention in the fMRI literature in recent years, where high amounts of serial correlation are inherent in the data structure. Prior to running ARMA models, we computed aggregated nodal time series by taking the average of all time series for voxels included in a given node, excluding those voxels that were missing or had no variance.

ARMA models represent the temporal dependence of observations in a time series, allowing one to remove the autoregressive components of the signal to achieve a “white” error time series. ARMA models were fit to each average nodal time series using the *Arima* function included in the *forecast* package in R (13) such that

$$\eta_t = c + \varepsilon_t + \sum_{i=1}^{p} \varphi_i \eta_{t-i} + \sum_{i=1}^{q} \theta_i \varepsilon_{t-i}$$

where $\varphi_i$ ... $\varphi_p$ denote freely estimated AR coefficients that quantify the degree of autocorrelation between the current realization ($\eta_t$) and previous realizations, and $\theta_i$ ... $\theta_p$ denote freely estimated MA coefficients that quantify the degree of dependence of the current innovation ($\varepsilon_t$) on prior innovations. The residual error term, also called the innovation term in an AR model, is assumed to be normally distributed (i.e. “white”) noise

$$\varepsilon_t \sim iid\ N(0, \sigma^2).$$

The *unique* coefficients of a given ARMA model for a node and subject act as a filter on the time series that whitens the residuals. However, the important quantity of interest in a graph theoretical analysis is the cross-correlation between every pair of nodal time series within a given subject, which are then used as cells in the adjacency matrix. Thus, to compute functional connectivity between regions, we stored the ARMA coefficients (sometimes referred to as the transfer function) for one node and used these to filter both time series for a given subject. We used the Pearson product-moment correlation coefficient to quantify the functional connectivity between nodal time series that had been passed through the same ARMA coefficients:

$$r_{\phi(y),\phi(x)} = \frac{1}{T-1}\sum_{t=1}^{T}\frac{\phi(y_t) - \overline{\phi(y)}}{s_{\phi(y)}} \cdot \frac{\phi(x_t) - \overline{\phi(x)}}{s_{\phi(x)}}$$

where $\phi(y)$ and $\phi(x)$ denote the time series for two nodes that have both been filtered by the fitted ARMA coefficients for $y$.

We fit a series of increasing complex ARMA models until the number of subject-wide "non-white" residuals fell below 5% of voxels for all subjects (i.e., the false positive rate on the test). Time series were deemed "non-white" on the basis of the Breusch-Godfrey test, which tested null hypotheses of serial correlation our nodal time series, which was computed up to six lags prior to the current realization (i.e., six seconds in the past). Through this procedure we retained our results from an ARMA(4,2) model as the edges of subject-level graphs for further analysis. We excluded one subject from all analyses who's residual time series remained non-white after ARMA(4,2) pre-whitening, whereas in all other subjects our prewhitening procedures were successful.

**Node selection: undirected analysis.** As described in the main text, our ultimate aim in the first step of our analysis was to cull those nodes in which no significant group differences in EC were likely to be detected to aid the reliability of CS-GIMME estimation. Thus, we sought to investigate which edge values had significant group differences in FC or had significant group x age interactions. Before conducting formal analyses, we aimed to determine if age-related changes in an edge's FC value conformed better to linear, inverse linear (asymptotic), or quadratic (u-shaped) functions of age. We first fit a series of regression models predicting edge FC as a function of subjects' mean FC values and interacting group and age variables. In order to test for different shapes of age-related effects we fit three regression models for each edge (i.e. every combination of nodes in the selected set), each corresponding to linear, inverse linear, and quadratic age effects (quadratic models also included a linear component). We ran this procedure for every edge and estimated linear, inverse linear, and quadratic age effects. We assumed a linear age effect to be the default but retained quadratic or inverse age effects for further analyses if a Vuong likelihood ratio test rejected the null hypothesis ($p < .05$) that the two models are equally close to the data generating process (14). Results indicated that eight edges were best fit by an inverse age-related parameterization, one was best fit by a quadratic function, and the rest fit by a simple linear trend. All significant age x group effects in our data were best fit by a linear trend.

The primary aim of our initial node selection analyses was to examine which edges best described age and BPD-related differences in FC. Given that our trimmed 19 x 19 adjacency matrix (after thresholding) would imply 154 separate tests, we sought to avoid running multiple separate models given that this mass univariate approach would have a high risk of false positive findings. Moreover, univariate analyses of edges would not provide insight into which subset of edges *jointly* discriminate FC differences as a function of BPD and age. With this in mind, a logistic ridge regression in which we predicted group status (BPD vs. HC) as a function of edge FC values for all edges in the joint set, as well as age-by-FC interactions from the age models selected above. Thus, the results we interpret are based on parameter estimates that jointly best discriminated between adolescents in the BPD and HC groups. By including edge x age interactions, the models were also able to identify nodes whose pattern of age-related change differed by group. This model was specified as

$$logit(BPD) = \beta_0 + \beta_1 Age + \beta_2 Age^2 + \beta_3 FC_{mean} + \beta_4 FC_{56-84} + \beta_5 FC_{56-84} \times Age + \cdots$$
$$+ \beta_{310} FC_{191-202} + \beta_{311} FC_{191-202} \times Age + \beta_{312} FC_{191-202} \times Age^2 + e$$

where $FC_{191-202}$ denotes the FC value between node 191 (R vmPFC) and node 202 (R BLA). In this example, the Age$^2$ term for $FC_{191-202}$ denotes that a quadratic age model fit better than linear or inverse variants according to the Vuong test, whereas for $FC_{56-84}$ the linear model fit the best.

We elected to use a regularized regression approach to overcome the p ≫ n problem in our model (hundreds of parameters, 82 observations), and handle high levels of collinearity in our data (15). Ridge regression shrinks model coefficients towards zero by penalizing the summed parameter estimates. This is achieved by augmenting the standard OLS loss function with an L2 penalty such that

$$L_{ridge}(\hat{\beta}) = \sum_{i=1}^{n}(y_i - x_i'\hat{\beta})^2 + \lambda\sum_{j=1}^{m}\hat{\beta}_j^2$$

where λ is a penalty parameter corresponding to the level of shrinkage on the standard OLS regression parameter estimate. The penalty parameter for our ridge regression was chosen using an automated selection algorithm implemented in the R *ridge* package(16,17). We elected to retain parameters as significant if their ridge *p*-value was < .01. We note that in such an analysis, there is no inherent need to correct for multiple comparisons since all edges are tested simultaneously. However, given the large number of potential contributing parameters in the fitted models we chose to retain a subset of results that were the most potent in distinguishing our groups and thus elected p < .01 as a more stringent test of significance.

**Effective connectivity network estimation and relations to group status.** GIMME estimates both lagged and contemporaneous relationships between nodal time series (and within time series, corresponding to AR processes) and benefits from relaxing the assumption that all individuals must be fit to the same model, yet utilizes regularities in individual subjects' path estimates to derive group-level edges that are estimated for every subject in the sample. Likewise, CS-GIMME extends upon this framework by also searching for edges that uniquely exist in one a priori subgroup (i.e. a clinical disorder, treatment group, neurostimulation condition, etc.) but not for another, thus allowing for the investigation of subgroup differences between edges that are estimated for all subjects (group-level edges) but also for the existence of edges that are unique to an a priori subgroup (18). We fit the CS-GIMME model to the preprocessed[1] time series using the *gimme* R package (19). Importantly, CS-GIMME authors have documented that in simulated data with no subgroup-specific edges, the algorithm simply estimates group and individual edges, effectively reducing the algorithm to the base GIMME approach.

**Directed nodal centrality estimation.** After obtaining parameter estimates for group-level, subgroup-level, and individual edges, we calculated in- and out-degree centrality for nodes that had either incoming or outgoing directed edges at the *group* (i.e. whole-sample) or *subgroup* (i.e. unique to BPD or controls) level: $k_i^{in} = \sum_{j \in N} w_{ji}$ and $k_i^{out} = \sum_{j \in N} w_{ij}$ (20,21). In the current case, $w_{ji}$ and $w_{ij}$ correspond to the beta estimate (extracted from CS-GIMME) for an edge traveling from node j to node i or from node i to node j, respectively. Centrality calculations were conducted using the *igraph package* in R (22). Importantly, in- and out-degree[2] were calculated only using contemporaneous paths from CS-GIMME, as we had no a priori suspicions

[1] While time series were preprocessed before EC estimation, we used the non-white nodal time series, rather than fitting CS-GIMME to a signal that had already been passed through an ARMA filter, given that GIMME explicitly contains an AR component.

[2] In formal graph theory, $k_i^{in}$ and $k_i^{out}$ are referred to as in- and out-*strength*, given that they consider directed edge weight, with in- and out-*degree* referring to a special (binary) case of strength. However, given the widespread use of degree centrality in neuroscience, we elect to keep our terminology close to the literature.

that lagged associations would benefit estimations of directed degree and is viewed as modeled nuisance variation. Further, in BOLD fMRI data the slow time course of hemodynamic activity likely implies that contemporaneous associations may reflect temporal causality even though strictly speaking GIMME estimates the directed influence of one node on another within the same TR.

In-degree and out-degree are graph theoretic metrics of directed centrality that denote in our case the summed beta coefficients of edges entering and exiting a node respectively. Thus, we were able to quantify which nodes play larger roles in the overall flow of information in the fronto-limbic circuit. While we only estimated directed degree for those nodes that had incoming or outgoing edges estimated by CS-GIMME at the *group* and *subgroup* level, these estimates contain information that is unique for each subject and thus contain information about individual-specific and subgroup-level edges. Although subgroup-level edges are estimated for every subject in a specific subgroup, a few subjects in the other group also tended to have that edge estimated at the individual level (typically 3-6 subjects in the opposing group had individual edges which were estimated for the entire other subgroup), demonstrating GIMME's ability to balance group regularities with the unique properties of individuals. In other words, we computed directed centrality estimates for nodes with incoming and outgoing edges at the group/subgroup level, though these estimates at the single-subject level reflect the sum of group, subgroup and individual edges.

Similar to our node selection step with undirected edges from above, we were interested in the ability of directed edges estimated at the group level in CS-GIMME to discriminate groups based on edge values or edge x age interactions. Thus, we fit two logistic multiple regression models predicting group status by directed edges and directed degree estimates in addition to age interactions in order to specify higher or lower connectivity in the BPD or control group or if directed connectivity (at the edge level or directed nodal centrality level) showed differential age-related connectivity changes between groups. In this case, regularization was not required, given the low levels of collinearity between directed edges (see Fig S2), which was not the case when considering undirected FC.

**Preliminary centrality regression models.** After estimating directed centrality estimates, before fitting the joint model described in the main text we preliminarily tested for group differences in directed centrality (in- and out-degree) by fitting a series of regression models regressing directed centrality estimates on group, age, and their interaction:

$$k_i^{in/out} = \beta_0 + \beta_1 Group + \beta_2 Age + \beta_3 Group\ x\ Age + e$$

where $k_i^{in/out}$ represents directed centrality estimates for node $i$ and Group is a binary predictor denoting group status (BPD = 1). Results from these regressions are summarized in Table S2.

**Preliminary UPPS mixed effects regression models.** Prior to fitting UPPS scales in the LGCM framework, we tested the ability of group status, age, and their interaction to predict impulsive symptoms in a series of mixed effects regression models (23):

$$L1{:}\ UPPS_j = \beta_{0j} + \beta_1 Group + \beta_2 Age + \beta_3 Group\ x\ Age + r_{ij}$$

$$L2{:}\ \beta_{0j} = \gamma_{00} + u_{0j}$$

where $\beta_{0j}$ reflects a linear combination of a group-level and subject-specific intercept terms. A separate model was fit for each of the five scales of the UPPS.

### Supplemental Results

**Node selection: undirected analysis.** Results from the logistic ridge regression are displayed in Table S1. We found significant group x age interactions between the right VS and the left ventromedial (t = -3.21, p = 0.001), rostromedial (t = -3.21, p = 0.001), and dorsomedial (t = -2.59, p = 0.01) prefrontal cortex. Adolescents with BPD symptoms decreased in mPFC-VS FC over time whereas adolescents in the control group increased in FC over development. Functional connectivity between left VS and left dACC was significantly lower in the BPD group (t = -2.78, p = 0.005) and that FC between left CeN and the bilateral dmPFC ($t_{right}$ = -2.90, p =0.004; $t_{left}$ = -2.59, p = .01) was significantly lower in the BPD group. Conversely, FC between the right BLA and right rmPFC was significantly higher in the BPD group (t = 2.58, p = .01).

This initial node selection analysis indicated that at the level of undirected FC, multiple nodes were not implicated in group status (Table 2, S1). We excluded 7 nodes on this basis, largely from the right mPFC/ACC and retained timeseries from 12 nodes for CS-GIMME estimation. While our undirected analysis did not show evidence of altered FC in two of the four amygdala nodes, we elected to retain all subcortical nodes for EC estimation due to a priori interest in these regions.

**Group-level edge analysis.** At the level of overall differences in group-level edges between participants in the BPD and control groups, we found modest support for greater edge strength in an edge from the left CeN to the right CeN (t = -1.98, p = .051) in the healthy control group. Age-related and age-modulated-by-group effects on group-level edges were all nonsignificant (all p's ≥ .108). Additional exploratory analyses of age-related changes in subgroup-level edges were also nonsignificant (all p's ≥ .398).

**Table S1. Results of ridge regression on undirected edge values**

| Edge (CS-GIMME labels) | Coefficient | Est.(S.E.) | t-score |
|---|---|---|---|
| L vmPFC – R VS (1-12) | Group x Age | -0.011(.004) | -3.21** |
| L rmPFC – R VS (2-12) | Group x Age | -0.011(.003) | -3.21** |
| L dmPFC – R VS (3-12) | Group x Age | -0.009(.004) | -2.59* |
| L dmPFC – L CeN (3-9) | Group | -0.009(.004) | -2.59* |
| L dACC – L VS (4-11) | Group | -0.010(.004) | -2.78* |
| R rmPFC – R BLA (5-8) | Group | 0.009(.004) | 2.58* |
| R dmPFC – L CeN (6-9) | Group | -0.010(.004) | -2.90** |

*Note.* Describes the set of edges that jointly discriminated groups in a logistic regression. Each row represents a coefficient in the model that significantly predicted group status at the $p < .01$ level (*$p < .01$, **$p < .005$.). For group effects, negative t-scores indicate that this edge was significantly lower in the BPD group compared to controls.

**Table S2. Nodal centrality metrics: unconditional relations to group, age, and their interaction**

| Node (CS-GIMME label) | Centrality | Coefficient | Est.(S.E.) | t-score (p value) |
|---|---|---|---|---|
| L vmPFC (1) | In | Group | -0.312(.222) | -1.40 (.16) |
| | | Age | -0.031(.161) | -0.19 (.85) |
| | | Group x Age | -0.006(.224) | -0.03 (.98) |
| | Out | Group | -0.029(.220) | -0.12 (.90) |
| | | Age | 0.142(.159) | 0.89 (.38) |
| | | Group x Age | 0.122(.221) | 0.55 (.58) |
| **L rmPFC (2)** | **In** | **Group** | **0.433(.220)** | **1.97 (.05)**[†] |
| | | Age | -0.008(.159) | -0.05 (.96) |
| | | Group x Age | -0.046(.221) | -0.21 (.84) |
| | Out | Group | | |
| | | Age | | |
| | | Group x Age | | |
| L dmPFC (3) | In | Group | | |
| | | Age | | |
| | | Group x Age | | |
| | Out | Group | 0.162(.222) | 0.73 (.47) |
| | | Age | -0.166(.161) | -1.03 (.31) |
| | | Group x Age | 0.060(.224) | 0.27 (.79) |
| **L dACC (4)** | In | Group | -0.144(.223) | -0.65 (.52) |
| | | Age | -0.148(.161) | -0.91 (.36) |
| | | Group x Age | 0.058(.224) | 0.26 (.80) |
| | **Out** | **Group** | **-0.936(.194)** | **-4.81 (<.001)**[***] |
| | | Age | -0.061(.141) | -0.43 (.67) |
| | | Group x Age | -0.173(.196) | -0.88 (.38) |
| **R rmPFC (5)** | **In** | **Group** | **0.515(.214)** | **2.41 (.02)**[*] |
| | | Age | 0.002(.155) | 0.01 (.99) |
| | | Group x Age | -0.264(.215) | -1.23 (.22) |
| | Out | Group | 0.168(.221) | 0.76 (.45) |
| | | Age | 0.215(.160) | 1.34 (.19) |
| | | Group x Age | -0.117(.223) | -0.92 (.60) |
| R dmPFC (6) | In | Group | 0.081(.224) | 0.36 (.72) |
| | | Age | 0.002(.162) | 0.015 (.99) |
| | | Group x Age | 0.151(.225) | 0.67 (.50) |
| | Out | Group | 0.035(.220) | 0.161 (.87) |
| | | Age | 0.025(.160) | 0.16 (.88) |
| | | Group x Age | -0.313(.222) | -1.41 (.16) |
| **L BLA (7)** | **In** | **Group** | **-0.443(.219)** | **-2.02 (.05)**[*] |

| | | | | |
|---|---|---|---|---|
| | | Age | 0.050(.159) | 0.314 (.76) |
| | | Group x Age | -0.117(.221) | -0.53 (.60) |
| | **Out** | **Group** | **0.581(.215)** | **2.70 (<.01)** ** |
| | | Age | 0.052(.156) | 0.33 (.74) |
| | | Group x Age | -0.120(.216) | -0.56 (.58) |
| **R BLA (8)** | **In** | **Group** | **-0.671(.208)** | **-3.23 (<.01)** ** |
| | | Age | -0.193(.151) | -1.28 (.20) |
| | | **Group x Age** | **0.359(.209)** | **1.72 (.09)** † |
| | **Out** | **Group** | **0.583(.221)** | **2.77 (<.01)** ** |
| | | Age | 0.237(.153) | 1.56 (.12) |
| | | **Group x Age** | **-0.392(.212)** | **-1.85 (.07)** † |
| **L CeN (9)** | In | Group | -0.011(.224) | -0.05 (.96) |
| | | Age | 0.098(.162) | 0.60 (.55) |
| | | Group x Age | -0.191(.226) | -0.85 (.40) |
| | **Out** | **Group** | **-0.468(.213)** | **-2.20 (.03)** * |
| | | **Age** | **-0.317(.154)** | **-2.06 (.04)** * |
| | | **Group x Age** | **0.359(.214)** | **1.67 (.10)** † |
| **R CeN (10)** | **In** | **Group** | **0.783(.196)** | **3.99 (<.001)** *** |
| | | Age | -0.088(.142) | -0.62 (.54) |
| | | Group x Age | -0.322(.198) | -1.63 (.11) |
| | **Out** | **Group** | **-1.162(.183)** | **-6.36 (<.001)** *** |
| | | Age | 0.062(.132) | 0.47(.64) |
| | | Group x Age | -0.026(.184) | -0.14 (.89) |
| **L VS (11)** | **In** | **Group** | **-0.625(.212)** | **-2.95 (<.01)** ** |
| | | Age | 0.183(.154) | 1.19(.24) |
| | | Group x Age | -0.159(213) | -0.74 (.46) |
| | Out | Group | 0.002(.222) | 0.01 (.99) |
| | | Age | 0.031(.161) | 0.20 (.85) |
| | | Group x Age | -0.249(.224) | -1.11 (.27) |
| R VS (12) | In | Group | -0.041 (.224) | -0.18 (.86) |
| | | Age | 0.135(.162) | 0.832 (.41) |
| | | Group x Age | -0.204(.225) | -0.91 (.37) |
| | Out | Group | | |
| | | Age | | |
| | | Group x Age | | |

**Table S3. Fixed effects of group, age, and their interaction on UPPS scales**

| UPPS scale | Effect | Est.(S.E.) | t-score |
|---|---|---|---|
| **Negative Urgency** | **Age** | **-0.044(.017)** | **-2.568*** |
| | Group | -0.036(.503) | -0.071 |
| | **Group x Age** | **0.053 (.024)** | **2.243*** |
| Premeditation (Lack of) | Age | -0.019(.015) | -1.247 |
| | Group | 0.320(.0444) | 0.719 |
| | Group x Age | 0.005(.021) | 0.224 |
| Perseverance (Lack of) | Age | -0.018(.014) | -1.308 |
| | Group | 0.193(.408) | 0.474 |
| | Group x Age | 0.024(.019) | 1.245 |
| Sensation Seeking | Age | -0.023(.022) | -1.016 |
| | Group | -0.584(.658) | -0.887 |
| | Group x Age | 0.033(.031) | 1.060 |
| **Positive Urgency** | **Age** | **-0.048(.016)** | **-3.065***** |
| | Group | -0.454(.0461) | -0.985 |
| | **Group x Age** | **0.054(.022)** | **2.468*** |
| **UPPS Total Score** | **Age** | **-0.030(.012)** | **-2.469*** |
| | Group | -0.116(.362) | -0.322 |
| | **Group x Age** | **0.034(.017)** | **1.992*** |

*Note.* A separate model was fit iteratively for each subscale, with the syntax: `lmer(UPPS_scale ~ Group*Age + (1|Subject))` in lme4, given that individual subjects had up to three UPPS measurements reflecting non-independence of all UPPS scores (i.e. UPPS scores nested within subjects). Coefficient estimates and S.Es are fixed effects estimates, reflecting group-average effects which take into account subject-specific random intercepts. Visual depiction can be found in Fig 2. Significant effects are bolded with corresponding significance level (*p < .05, **p < .01, ***p < .005). To verify the often-reported group differences in impulsive symptoms, we ran two-sample t-tests on each of the UPPS subscales and confirmed strong group effects on all UPPS subscales besides sensation seeking, such that the BPD group was higher on level of impulsivity.

Figure S1.

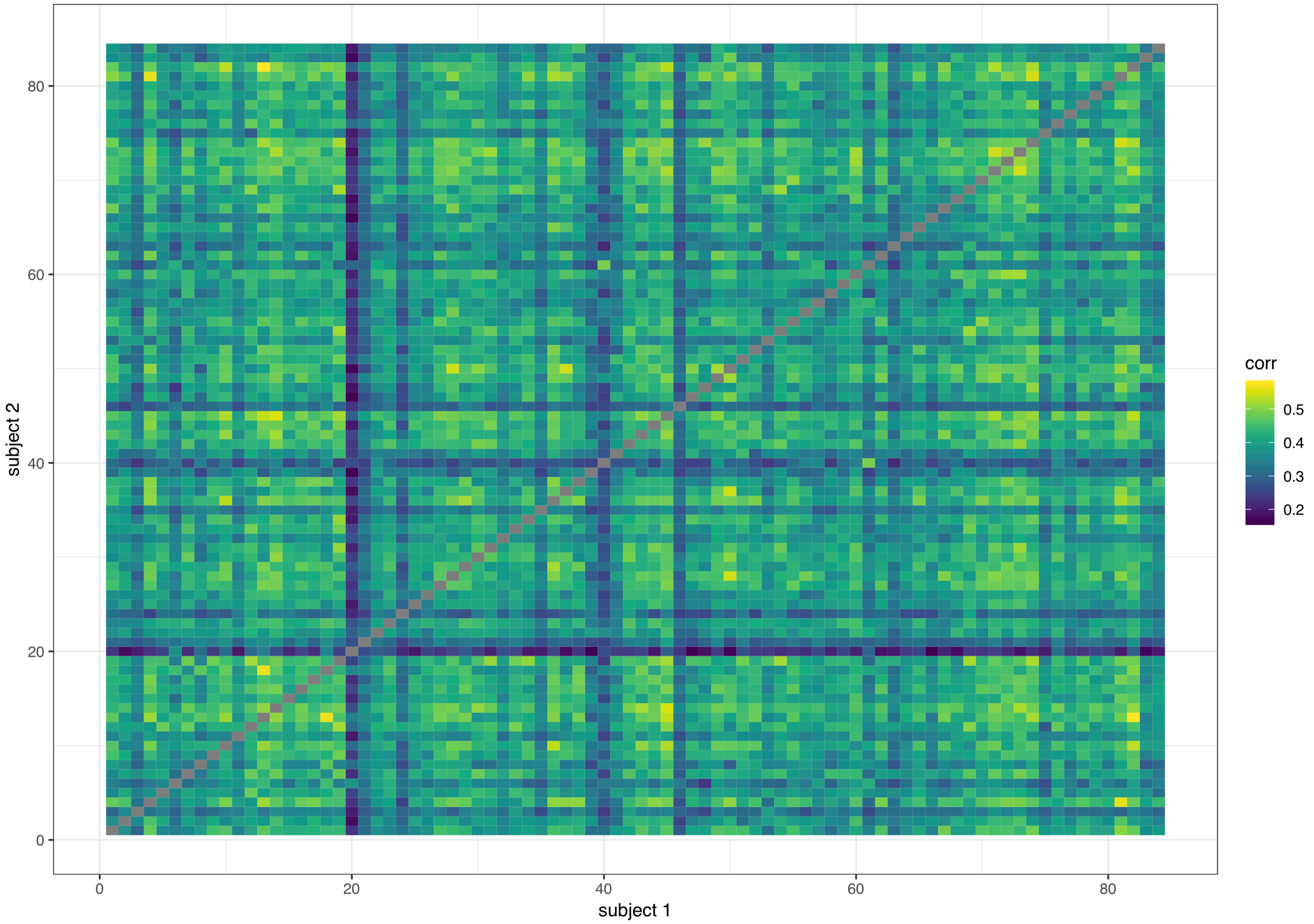


*Note.* Static pearson correlations of individual adjacency matrices by subject. This was conducted on the original 84 subjects that passed our excessive motion screen and is simply meant to demonstrate thinking regarding removal of one subject in the BPD group (who can be spotted by eye in row/column 20) whose adjacency matrix had an unusually low correlation on the whole (~.1) with all other subjects in the study and showed divergence measured by the Mahalanobis distance.

Figure S2.

*Correlation matrix: group, age, ERI measures, and amygdala edges and centrality estimates*

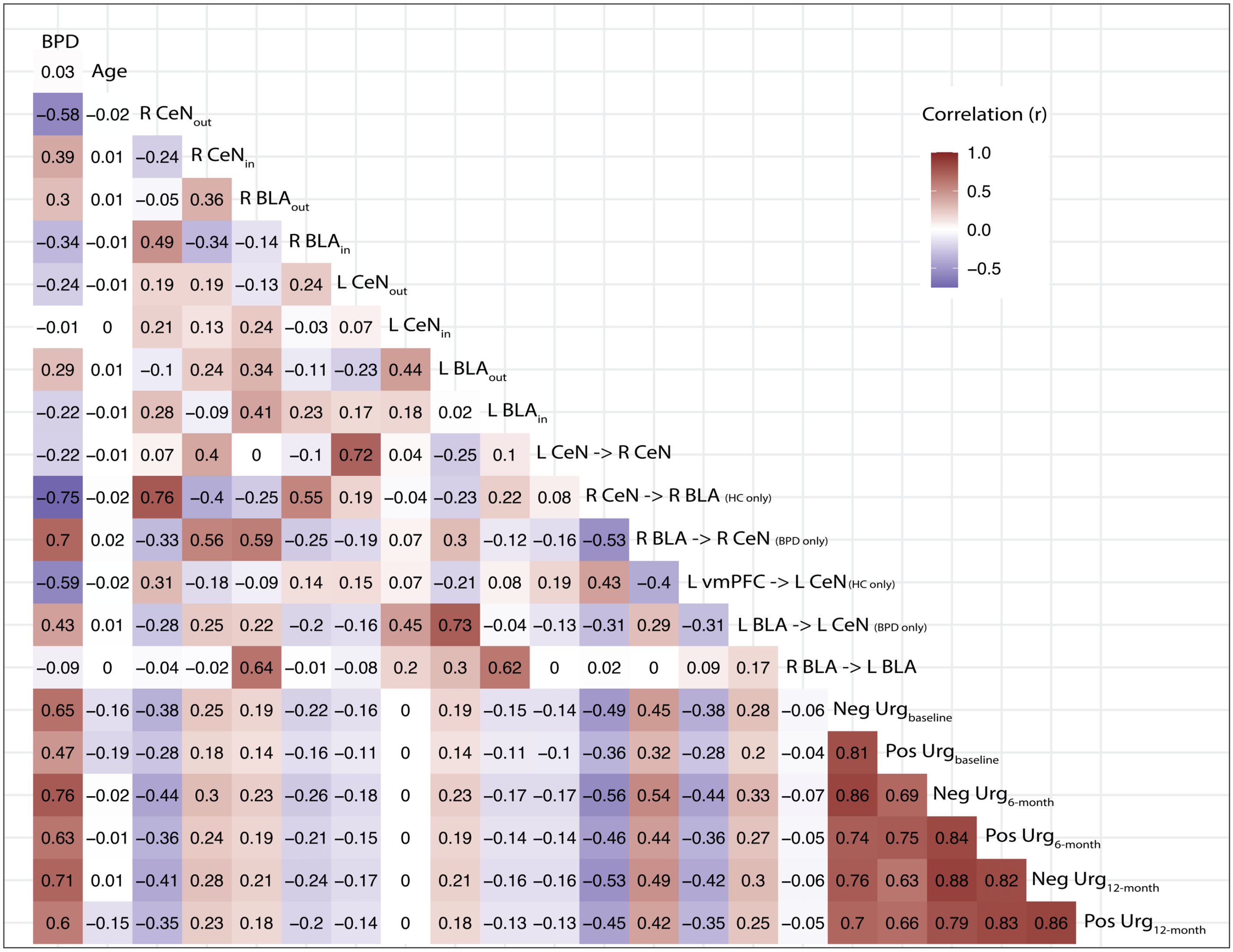


*Note.* "in" and "out" subscripts denote estimates of *nodal* in- and out- degree centrality, respectively. Labels with arrows denote beta estimates for a given *edge* estimated via CS-GIMME. Arrows denote the direction of influence from one node to another and edges with parenthetical designations "(HC only)", denote edges that were estimated at the subgroup level for all subjects in a given a priori subgroup.

Figure S3.

*Detailed directed edge counts*

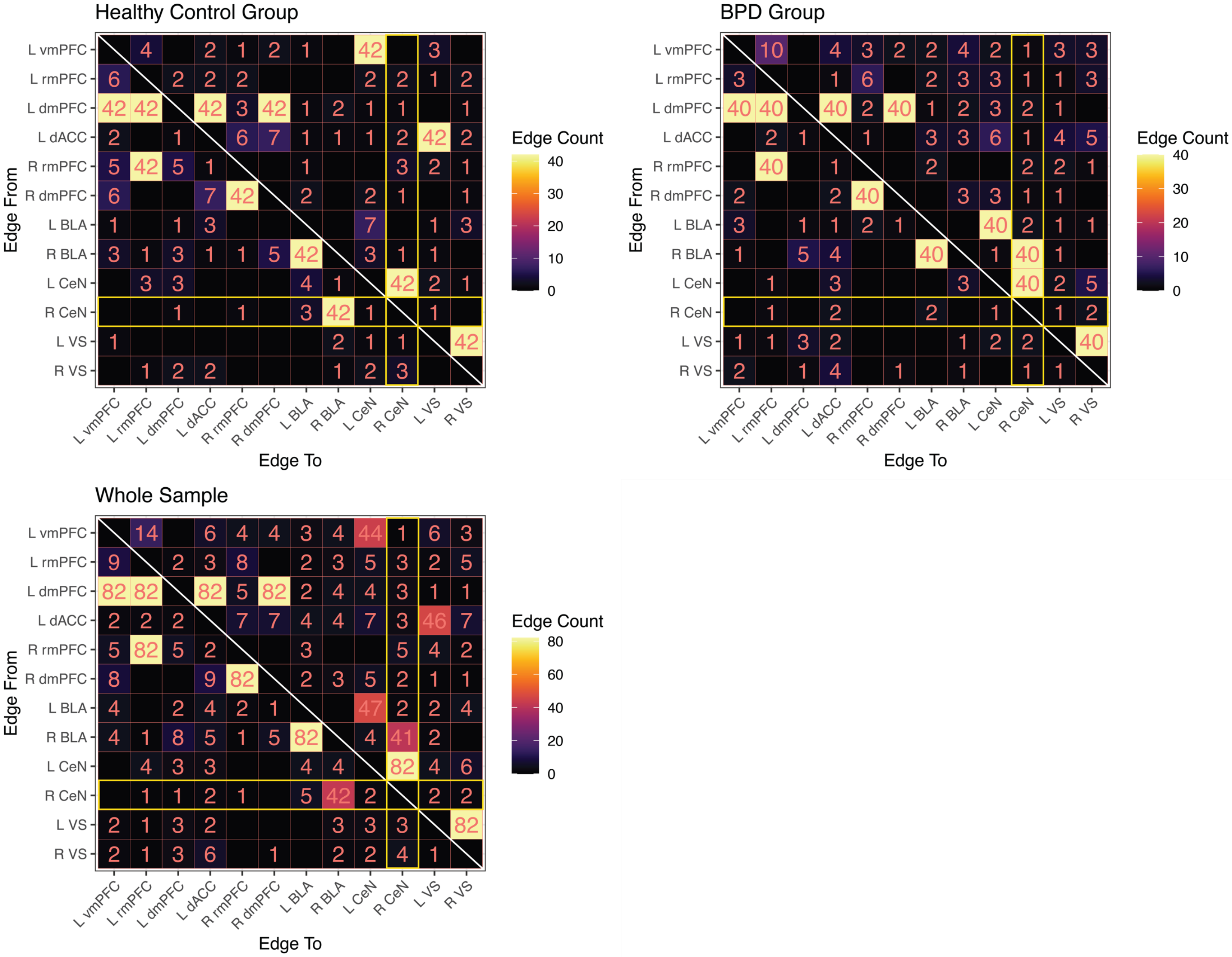


*Note.* To visualize the composition of nodal centrality metrics, we show the presence of specific edges estimated within the fronto-limbic circuit estimated from CS-GIMME. The top pane shows edge counts split between groups with the fill of each cell denoting the number of subjects with an edge estimated from the node on the y-axis to the node on the x-axis. Since centrality of R CeN was of central importance in the current study, we highlight these cells in yellow. Notice the dissociation in R BLA edges in subgroups, with sparse edges estimated at the individual levels for other nodes.
